# Myosin-X recruits lamellipodin to filopodia tips

**DOI:** 10.1101/2022.08.17.504298

**Authors:** Mitro Miihkinen, Ana Popović, Sujan Ghimire, Rafael Saup, Max L.B. Grönloh, Neil J. Ball, Benjamin T. Goult, Johanna Ivaska, Guillaume Jacquemet

**Author notes:** Equal contribution (Mitro Miihkinen, Ana Popović).

## Abstract

Myosin-X (MYO10), a molecular motor localizing to filopodia, is thought to transport various cargo to filopodia tips, modulating filopodia function. Yet, only a few MYO10 cargoes have been described. Here, using GFP-Trap and BioID approaches combined with mass spectrometry, we identified lamellipodin (RAPH1) as a novel MYO10 cargo. We report that the FERM domain of MYO10 is required for RAPH1 localization and accumulation at filopodia tips. Previous studies have mapped RAPH1 interaction with adhesome components to its talinbinding and Ras-association domains. Surprisingly, we find that the RAPH1 MYO10-binding site is not within these domains. Instead, it comprises an area with previously unknown functions. Functionally, RAPH1 supports MYO10 filopodia formation and stability but is not involved in regulating integrin activity in filopodia tips. Taken together, our data indicate a feed-forward mechanism whereby MYO10 filopodia are positively regulated by MYO10-mediated transport of RAPH1 to the filopodium tip.

## Introduction

Cell migration is essential during embryonic development, immune surveillance, and wound healing. Misregulation of cell migration is implicated in multiple diseases, including inflammation and cancer. One hallmark of cell motility is a high degree of plasticity, allowing cells to adopt different morphologies and migration modes (Conway and Jacquemet, 2019). A shared feature of efficient cell migration is the ability of cells to probe and interact dynamically with their environments using cellular protrusions such as filopodia, lamellipodia, or pseudopods.

Filopodia are small and dynamic finger-like actin-rich protrusions (1-5 mm in length and 50-200 nm in width) and are often the first point of contact between a cell and its immediate surroundings. Filopodia contain cell-surface receptors, such as integrins, cadherins, and growth factor receptors, interacting with and interpreting various extracellular cues. Filopodia assembly is primarily driven by the linear polymerization of actin filaments with their barbed ends facing the plasma membrane (Jacquemet et al., 2015). These filaments are further organized into tightly packed bundles by actin-bundling proteins. This unidirectional organization allows molecular motors, such as myosin-X (MYO10), to walk along filopodia and accumulate at their tips (at approximately 600 nm/s) (Kerber et al., 2009). By doing so, MYO10 is thought to transport various proteins to filopodia tips, modulating filopodia function (Jacquemet et al., 2015; Arjonen et al., 2014; Berg and Cheney, 2002; Hirano et al., 2011; Zhang et al., 2004). Yet, only very few MYO10 cargoes have been proposed to date, with the netrin DCC receptor (Zhu et al., 2007; Wei et al., 2011), integrins (Zhang et al., 2004; Wei et al., 2011) and VASP (Tokuo and Ikebe, 2004) being the principal ones. The MYO10 MyTH4/FERM domain (termed MYO10-FERM here for simplicity) domain has been described as the main cargo binding site in MYO10 (Wei et al., 2011). We previously reported that MYO10-FERM was not required to localize integrins or VASP at filopodia tips. Instead, we found that MYO10-FERM is required for proper integrin activation at filopodia tips (Miihkinen et al., 2021). In addition, we found that the deletion of the MYO10 FERM domain had little impact on the localization of significant filopodia tip complex components (Miihkinen et al., 2021). These results question the role of MYO10 as a cargo transporting molecule.

Here, we set out to identify novel MYO10 cargo molecules. Using GFP-Trap and BioID approaches combined with mass spectrometry, we identified RAPH1 as a novel MYO10-binding partner. Using structured illumination microscopy, we report that the MYO10’s FERM domain is required for RAPH1 localization and accumulation at filopodia tips; thus, RAPH1 is an MYO10 cargo. We map the RAPH1 MYO10-binding site to a previously uninvestigated RAPH1 sequence and demonstrate that RAPH1 is a critical positive regulator of filopodia formation and stability in cells. Our results indicate that, in filopodia, RAPH1 is not required for integrin activation. Instead, RAPH1 regulates MYO10 filopodia formation and stability.

## Results and discussion

### RAPH1 is a putative MYO10 cargo

To identify novel MYO10 cargo, we searched for proteins that interact specifically with MYO10-FERM, the main cargo-binding site in MYO10 (Wei et al., 2011). We performed GFP pull-downs in U2-OS cells stably expressing GFP, GFP-MYO10^FERM^, or GFP-Talin^FERM^ (talin-1 FERM domain), followed by mass spectrometry analysis (Fig. 1A and Fig. 1B). Talin^FERM^ was selected as an additional control as it shares structural similarities with the MYO10 FERM domain but performs different functions in cells (Miihkinen et al., 2021). We identified 87 proteins that were specifically enriched in the MYO10-FERM pull-downs (Fig 1A and 1B and Table S1). Interestingly, small GTPase regulators such as RASAL2, ARHG-DIA, or TRIO were among the enriched putative MYO10-FERM binders (Table S1).

**Fig. 1.**
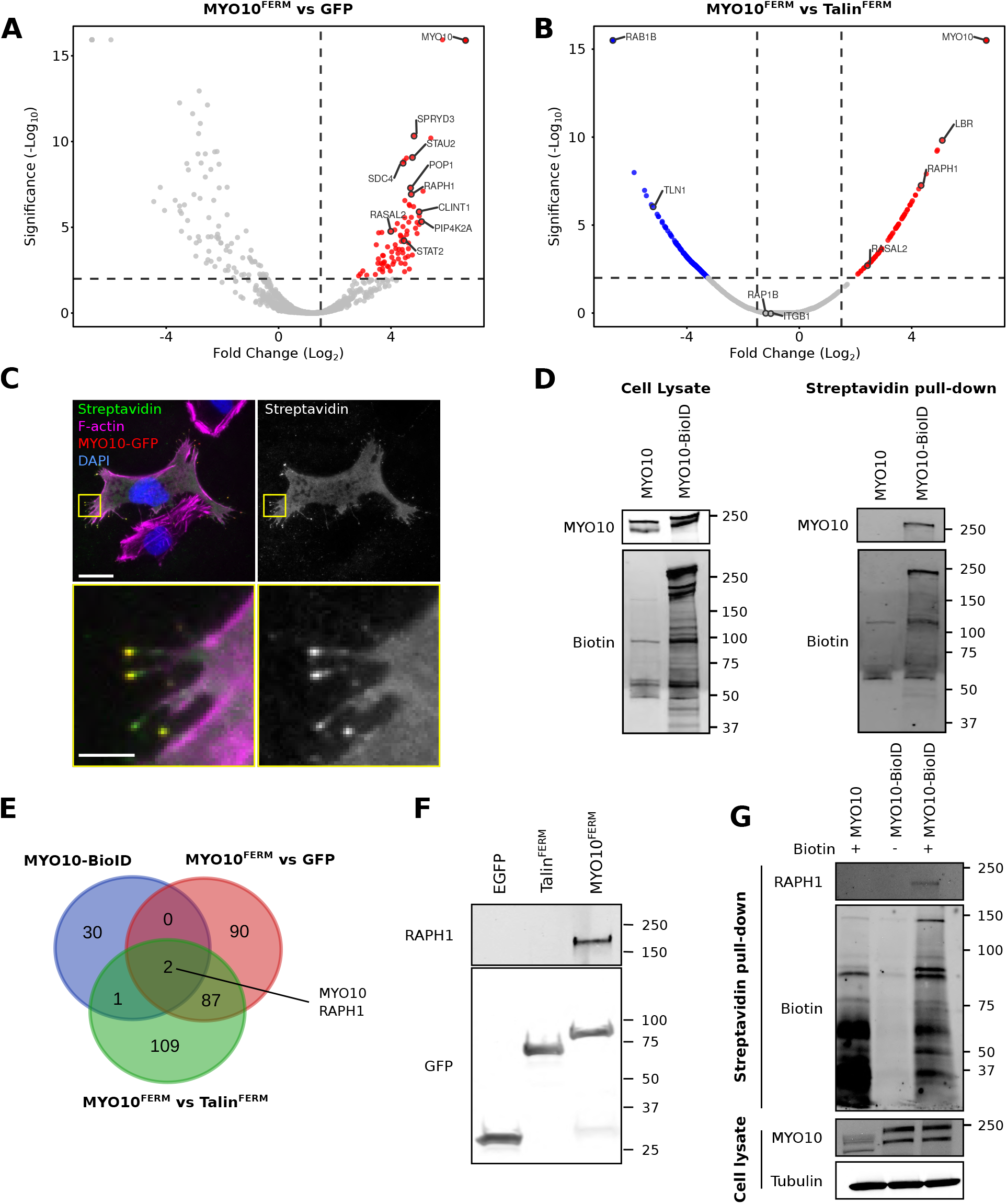
Mass-spectrometry analyses identify RAPH1 as a putative MYO10 binder. (**A**-**B**) Mass spectrometry (MS) analysis of GFP-MYO10^FERM^ and GFP-Talin^FERM^ binding proteins. Comparison of the GFP-MYO10^FERM^ dataset to GFP (**A**) and GFP-Talin^FERM^ (**B**) datasets are displayed as volcano plots where the fold-change enrichment is plotted against the significance of the association (see table S1 for the MS data). The volcano plots were generated using VolcaNoseR (Goedhart and Luijsterburg, 2020). (**C**) U2-OS cells transiently expressing GFP-MYO10-BioID were plated on fibronectin (FN) in the presence of biotin for 24 h, fixed, stained for biotinylated proteins (using streptavidin), F-actin, and Dapi, and imaged using a spinning disk confocal (SDC) microscope. The yellow squares highlight regions of interest (ROIs), which are magnified; scale bars: (main) 25 μm; (inset) 5 μm. Note that only one cell in this field of view expresses the GFP-MYO10-BioID construct. (**D**) U2-OS cells stably expressing GFP-MYO10-BioID or GFP-MYO10 were plated on FN for 24 h in the presence of biotin. Cells were then lysed, and biotinylated proteins purified using streptavidin beads. Recruited proteins were analyzed using western blot and MS (see Table S1 for the MS data). Western blots are displayed (representative of five biological repeats). (**E**) Venn diagram highlighting the overlap of MYO10-enriched proteins identified from the indicated MS datasets. (**F**) GFP pull-down in U2-OS cells expressing GFP-MYO10^FERM^, GFP-Talin^FERM^, or GFP alone. RAPH1 recruitment to the bait proteins was then assessed using western blot (representative of three biological repeats). (**G**) U2-OS cells stably expressing GFP-MYO10-BioID or GFP-MYO10 were plated on FN for 24 h in the presence or absence of biotin. Cells were then lysed and biotinylated protein purified using streptavidin beads. RAPH1 biotinylation was then assessed using western blots (representative of three biological repeats).

Next, to narrow the list of putative MYO10 cargo, we tagged GFP-MYO10 with the promiscuous biotin ligase BioID (Roux et al., 2012) adjacent to the MYO10 FERM domain. In cells, GFP-MYO10-BioID localized to and biotinylated proteins at filopodia tips (Fig. 1C). We purified biotinylated proteins in cells expressing GFP-MYO10 (negative control) or GFP-MYO10-BioID using streptavidin pulldowns (Fig. 1D) and performed mass spectrometry analyses. Somewhat unexpectedly, this approach identified very few proteins, perhaps due to the slow kinetics of the biotin ligase used (Fig. 1E and Table S2). Nevertheless, when comparing our GFP pull-down and BioID datasets, only two proteins, MYO10 itself and lamellipodin (RAPH1), were identified consistently as enriched to MYO10 over controls (Fig. 1E). Western blot analyses confirmed that RAPH1 co-purifies with GFP-MYO10^FERM^ and that RAPH1 is biotinylated in cells expressing GFP-MYO10-BioID (Fig 1F and 1G). These results led us to speculate that RAPH1 could be an MYO10 cargo.

### MYO10-FERM is required for RAPH1 localization at filopodia tips

RAPH1 is a member of the MRL (Mig-10/RIAM/Lamellipodin) protein family, with MIG-10 being the Caenorhabditis elegans ortholog of RAPH1 (Coló et al., 2012). RAPH1 was previously reported to localize to filopodia tips (Krause et al., 2004; Jacquemet et al., 2019), but its contribution to filopodia function remains unknown. Using structured illumination microscopy, we found that RAPH1 specifically accumulates at filopodia tips where it colocalizes with MYO10, while RIAM is uniformly distributed along filopodia (Fig. S1A). In addition, live-cell imaging at high spatiotemporal resolution indicated that RAPH1 closely follows MYO10 puncta at filopodia tips throughout the filopodia life cycle (Fig. S1B and Movie S1).

Our mass spectrometry data indicated that the MYO10 FERM domain recruits RAPH1. Therefore, we investigated the requirement for MYO10-FERM to localize RAPH1 to filopodia tips. We overexpressed an RFP-tagged MYO10 construct lacking the FERM domain (MYO10^ΔF^) in cells (Miihkinen et al., 2021), together with either RAPH1-GFP or VASP-GFP. Deleting the MYO10 FERM domain led to a loss of RAPH1 accumulation at filopodia tips (Fig. 2A-D), whereas VASP recruitment remained unaffected (Fig. 2A-D). In line with these results, the accumulation of endogenous RAPH1 was also lost at the tip of MYO10^ΔF^ filopodia (Fig. 2E-F). Taken together, these findings demonstrate that MYO10 and its FERM domain are required for RAPH1 accumulation at filopodia tips. These results also suggest that, despite containing multiple VASP-binding sites (Krause et al., 2004), RAPH1 is not a prerequisite for VASP localization to MYO10 filopodia.

**Fig. 2.**
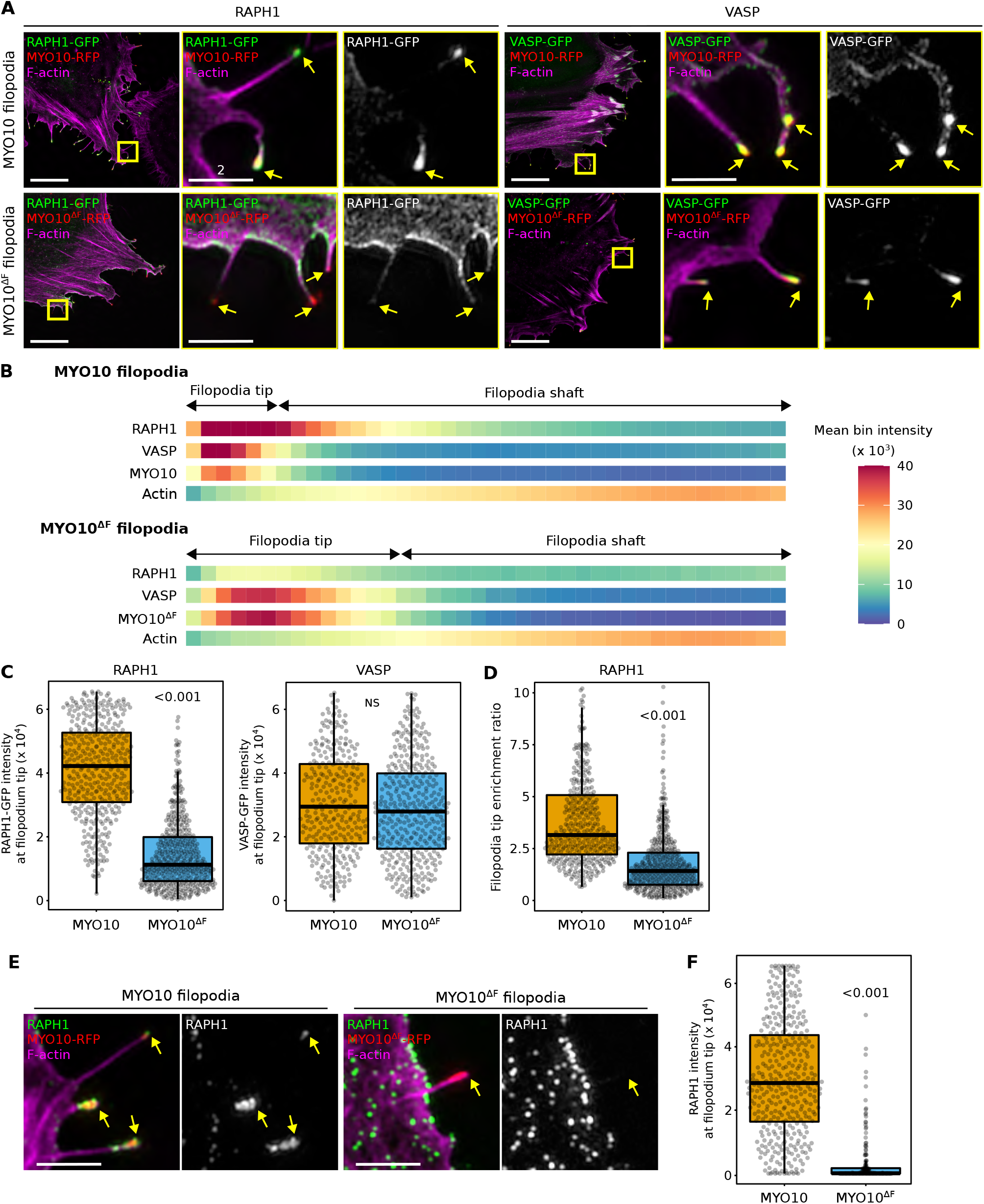
RAPH1 is recruited to filopodia tips in an MYO10-FERM-dependent manner. (**A**-**D**) U2OS cells expressing MYO10-RFP or MYO10^ΔF^-RFP together with RAPH1-GFP or VASP-GFP were plated on FN for 2 h, fixed, stained for F-actin, and imaged using SIM. (**A**) Representative MIPs are displayed. The yellow squares highlight ROIs, which are magnified; yellow arrows highlight filopodia tips; scale bars: (main) 20 μm; (inset) 2 μm. (**B**) Heatmap highlighting the sub-filopodial localization of the proteins imaged in (**A**) generated from intensity profiles (n > 300 filopodia per condition; three biological repeats). (**C**) The average intensity of RAPH1 and VASP staining at filopodia tips measured in (**B**) are displayed as box plots. (**D**) The preferential recruitment of RAPH1 to MYO10 or MYO10^ΔF^ filopodia tips over shafts was assessed by calculating an enrichment ratio (averaged intensity at filopodia tip versus shaft). Results are displayed as Tukey box plots. (**E**-**F**) U2-OS cells expressing MYO10-RFP or MYO10^ΔF^-RFP were plated on FN for 2 h, fixed, stained for F-actin and endogenous RAPH1, and imaged using SIM. (**E**) A representative ROI is displayed. Yellow arrows highlight filopodia tips; scale bars: 2 μm. (**F**) The average intensity of endogenous RAPH1 at filopodia tips is displayed as box plots (n > 175 filopodia per condition; three biological repeats). For all panels, p-values were determined using a randomization test. NS indicates no statistical difference between the mean values of the highlighted condition and the control.

### RAPH1 directly interacts with MYO10

Next, we sought to identify the MYO10-binding domain(s) within RAPH1. RAPH1 comprises several conserved domains, including a Ras-association (RA) and a pleckstrin homology (PH) domain. RAPH1 also contains known profilin-, VASP- and multiple putative SH3-binding sites (Fig. 3A). Furthermore, previous work indicated that RAPH1 binds to talin-FERM via two N-terminal talin-binding sites (Lee et al., 2009; Chang et al., 2014) (Fig. 3A and Fig. S2A). As talin-FERM and MYO10-FERM share structural similarities (Miihkinen et al., 2021), we speculated that RAPH1 could bind to MYO10-FERM via these talin-binding sites. To test this hypothesis, we generated a RAPH1 deletion construct lacking both talin-binding sites (Fig. S2B). Deleting both RAPH1 talin-binding sites did not affect RAPH1 localization to filopodia tips indicating that RAPH1 talin-binding sites are not required for MYO10 interaction (Fig. S2B).

**Fig. 3.**
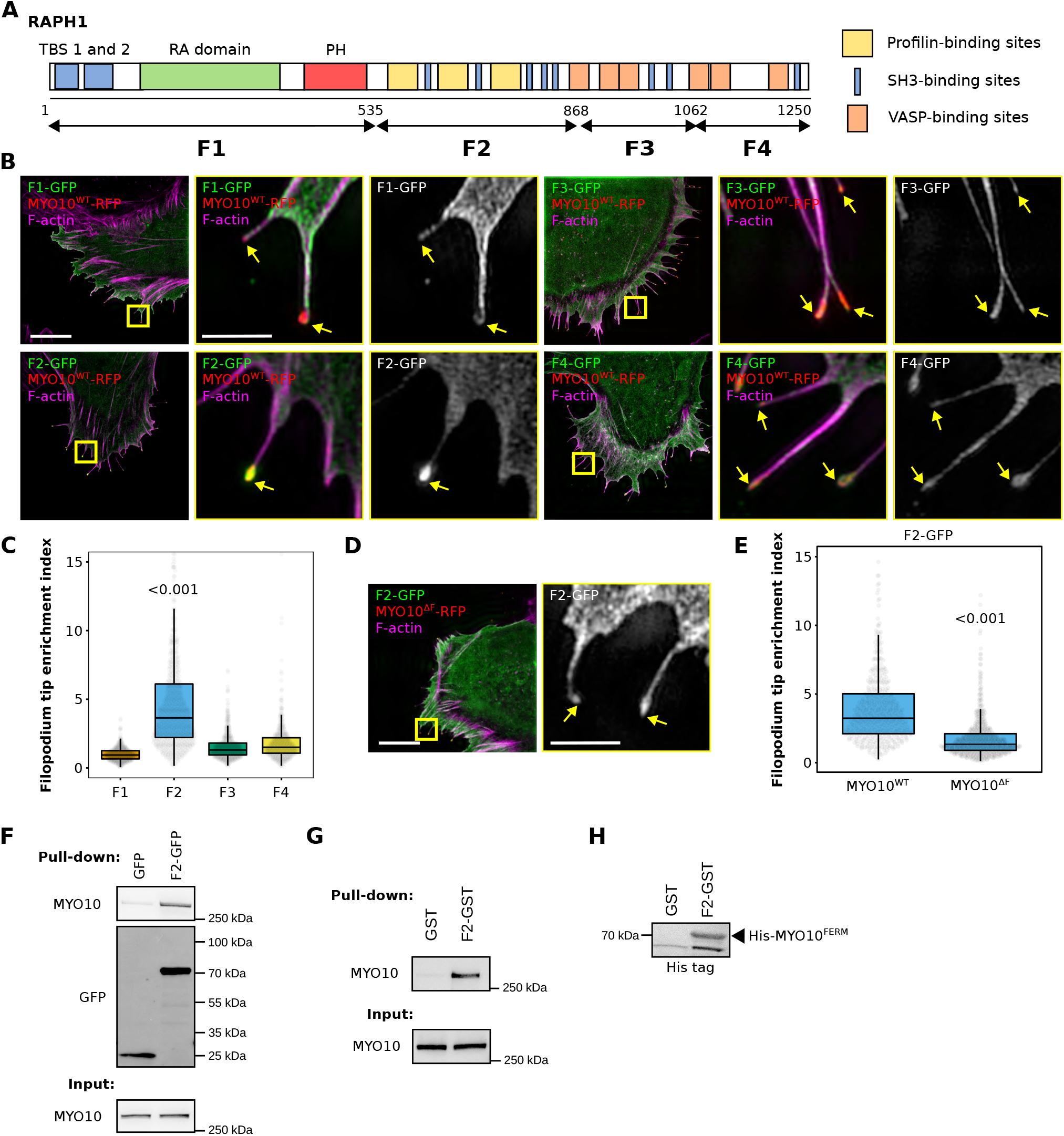
RAPH1 is recruited to filopodia tips via a previously uncharacterized region. (**A**) Cartoon representation of RAPH1 domains. The boundaries of the four fragments (F1 to F4) used in this study are highlighted. (**B**) U2-OS cells expressing MYO10-RFP and one of the four RAPH1 fragments were plated on FN for 2 h, fixed, stained for F-actin, and imaged using SIM. Representative MIPs are displayed. The yellow squares highlight ROIs, which are magnified; yellow arrows highlight filopodia tips; scale bars: (main) 10 μm; (inset) 2 μm. (**C**) The preferential recruitment of the four RAPH1 fragments to filopodia tips was assessed by calculating an enrichment ratio (averaged intensity at filopodia tip versus shaft; > 415 filopodia per condition, three biological repeats). (**D**) U2-OS cells expressing MYO10WT-RFP and the GFP-RAPH1^F2^ (GFP-F2) were plated on FN for 2 h, fixed, stained for F-actin, and imaged using SIM. Representative MIPs are displayed. The yellow squares highlight ROIs, which are magnified; yellow arrows highlight filopodia tips; scale bars: (main) 10 μm; (inset) 2 μm. (**E**) The preferential recruitment of GFP-RAPH1^F2^ to filopodia tips was assessed as in (**C**) (> 427 filopodia per condition, three biological repeats). (**F**) GFP pull-down in MDA-MB-231 cells expressing GFP-RAPH1^F2^, or GFP alone. Endogenous MYO10 recruitment to the bait proteins was then assessed using western blot (representative of three biological repeats). (**G**) Pull-down using recombinant GST-RAPH1^F2^ or GST alone in MDA-MB-231 cell lysates. MYO10 binding to GST-RAPH1^F2^ was then assessed using western blot (representative of three biological repeats). (**H**) Pull-down using recombinant GST-RAPH1^F2^ or GST alone and recombinant his-tagged MYO10^FERM^. MYO10^FERM^ binding to GST-RAPH1^F2^ was then assessed using western blot (representative of three biological repeats).

Next, we generated four truncated RAPH1 constructs (named F1 to F4; Fig. 3A) and mapped their filopodia localization (Fig. 3B). Somewhat surprisingly, the RAPH1 fragment F1 containing the PH and the RA domains did not accumulate at filopodia tips. Indeed, among the four constructs tested, only the RAPH1 F2 fragment, which contains the profilin binding sites, displayed an evident accumulation at filopodia tips (Fig. 3B and 3C). This required an intact MYO10 FERM domain and was lost in MYO10^ΔF^ filopodia (Fig. 3D and 3E), indicating that RAPH1 is recruited to filopodia tips via this F2 region.

To validate MYO10-RAPH1 binding, we performed GFP-trap experiments in MDA-MB-231 cells expressing GFP or GFP-RAPH1^F2^. We chose MDA-MB-231 cells for their high endogenous MYO10 protein levels (Jacquemet et al., 2019, 2016). MYO10 co-precipitated with GFP-RAPH1^F2^ (Fig. 3F), validating our microscopy-based assays. In addition, MYO10 was pulled down from cell lysates using recombinant GST-RAPH1^F2^ (Fig. S2C, Fig. 3G), and we detected binding between purified, recombinant GST-RAPH1^F2^ and his-tagged MYO10-FERM proteins (Fig. 3H).

Altogether, our data demonstrate a direct interaction between RAPH1 and MYO10 and that RAPH1-MYO10 binding is required for RAPH1 localization to filopodia tips. This interaction is mediated by the MYO10-FERM domain and a previously unexplored region within RAPH1 located after its PH domain.

### RAPH1 modulates filopodia formation and functions

Next, we investigated the contribution of RAPH1 to filopodia. RAPH1 silencing with two independent siRNA oligos in U2-OS cells expressing MYO10-GFP significantly reduced MYO10-positive filopodia numbers and filopodia length (Fig. 4A, 4B, and 4C). Interestingly, in a small proportion of RAPH1-silenced cells (below 1%), the filopodia tip complex collapsed, as observed by the dispersed localization of MYO10 along the filopodia shaft (Fig. 4C). While this phenotype was rare, it was not observed in control cells.

**Fig. 4.**
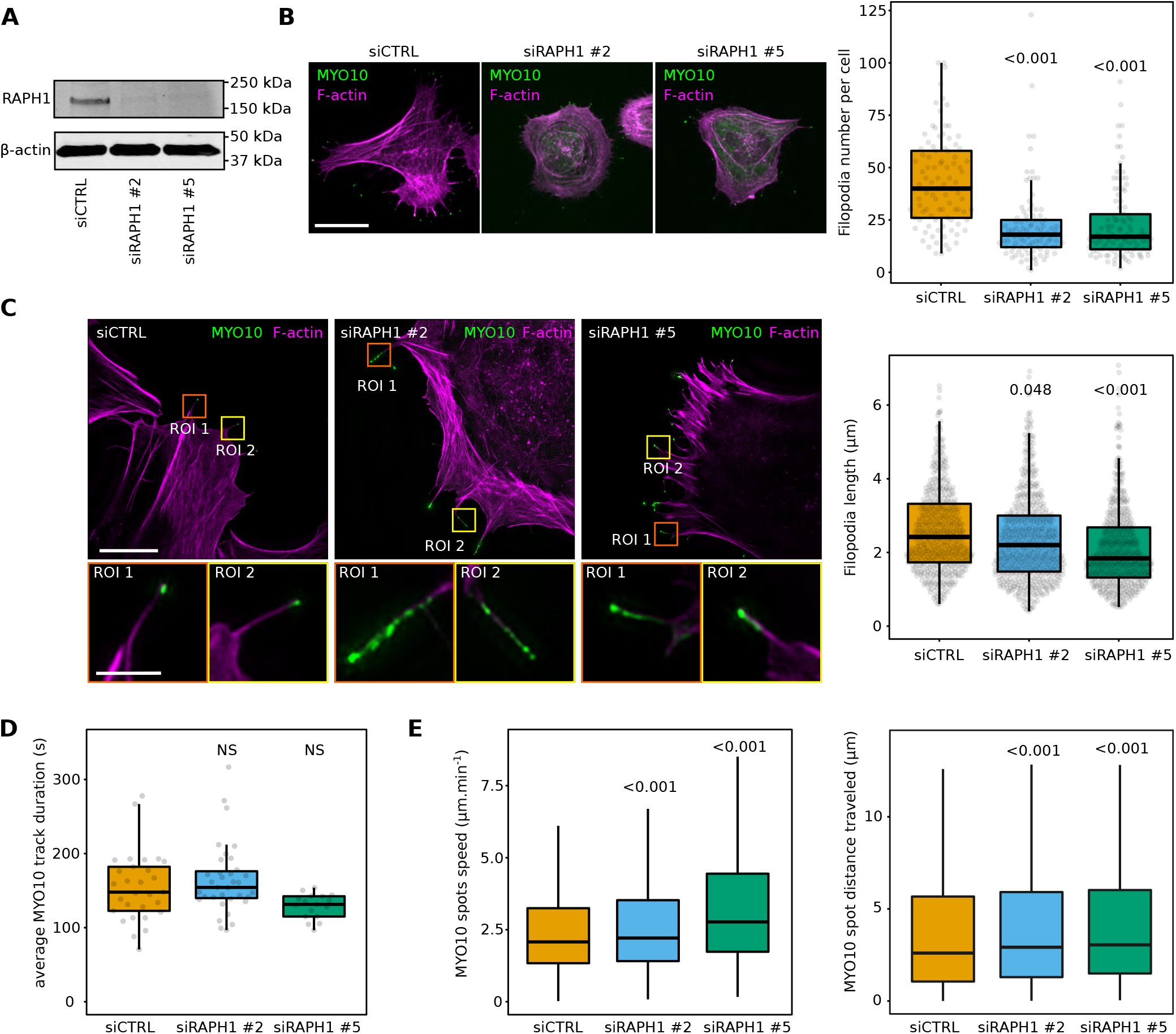
RAPH1 supports filopodia formation and the stability of the filopodia tip complex. (**A**) Efficiency of siRNA-mediated RAPH1 silencing using two different siRNA oligos in U2-OS cells. A representative western blot is displayed. (**B**) RAPH1-silenced U2-OS cells transiently expressing MYO10-GFP were plated on FN for 2 h, fixed, and the number of MYO10-positive filopodia per cell was quantified (n > 93 cells per condition, three biological repeats). (**C**) RAPH1-silenced U2-OS cells transiently expressing MYO10-GFP were plated on FN, stained for F-actin and imaged using SIM. Representative MIPs are displayed; scale bars: (main) 20 μm; (inset) 2 μm. Quantifications of filopodia length from SIM images are displayed (n > 530 filopodia per condition; three biological repeats). (**D**-**E**) RAPH1-silenced U2-OS cells transiently expressing MYO10-GFP were plated on FN and imaged live using an Airyscan confocal microscope (1 picture every 5 s over 20 min). For each condition, MYO10-positive particles were automatically tracked. (**D**) The average MYO10 track duration per cell is displayed (three biological repeats, n > 17 cells per condition). (**E**) the average speed and the total distance traveled by MYO10 spots are displayed (n > 9600 filopodia; three biological repeats). For all panels, p-values were determined using a randomization test. NS indicates no statistical difference between the mean values of the highlighted condition and the control.

RIAM and RAPH1 have been implicated in modulating integrin activity (Lee et al., 2009). While this role is now well established for RIAM, it is more controversial for RAPH1 (Coló et al., 2012; Chang et al., 2014; Lafuente et al., 2004; Watanabe et al., 2008). As the MYO10 FERM domain is required to activate integrin at filopodia tips (Miihkinen et al., 2021), we next investigated the role of RAPH1 in modulating integrin activity at filopodia tips (Fig. S3). Using SIM and filopodia mapping analyses, we found that integrin activation at filopodia tip is comparable or slightly elevated in RAPH1-depleted cells compared to CTRL, indicating that RAPH1 is not required for integrin activation at filopodia tips (Fig. S3).

Finally, we explored the role of RAPH1 in modulating filopodia dynamics in control and RAPH1-silenced U2-OS cells expressing MYO10-GFP. While the overall filopodia lifetime was unaffected after RAPH1 depletion, MYO10 puncta moved faster and over longer distances than in control cells (Fig. 4D and 4E), indicating that the filopodia tip complex is more dynamic in RAPH1 depleted cells.

Our results demonstrate that (1) MYO10 is required to target RAPH1 to filopodia tips and (2) RAPH1 contributes to MYO10 filopodia formation and dynamics. We propose that MYO10 transports RAPH1 to filopodia tips contributing to filopodia stability via yet unknown mechanisms, possibly involving RAPH1 interactions with other proteins such as VASP. However, our data do not exclude the possibility that RAPH1 simply diffuses to filopodia and that MYO10 only contributes to RAPH1 accumulation at filopodia tips without direct transport. Testing this would require performing two-color, single-molecule imaging of MYO10 and RAPH1 to see if these proteins move toward filopodia tips together. However, we find that RAPH1 is not very abundant in filopodia when the MYO10 FERM domain is missing, suggesting that RAPH1 is likely to be actively transported by MYO10.

At filopodia tips, RAPH1 is presumably in a complex with MYO10, VASP, and actin. In this scenario, MYO10 could tether RAPH1 to filopodia tips using its motor domain, providing resistance against the retrograde actin flow in filopodia (Bornschlögl, 2013; Lidke et al., 2005). Once tethered, RAPH1 would cluster and increase VASP activity by tethering VASP to the actin filaments (Hansen and Mullins, 2015). While we found that VASP molecules still localize to filopodia tips in the absence of RAPH1, VASP activity will likely be reduced, which could explain the shorter filopodia observed in RAPH1-silenced cells. Therefore, we propose that MYO10-mediated transport of RAPH1 to the filopodium tip is a feed-forward mechanism that positively regulates MYO10 filopodia.

Interestingly, both MYO10 and RAPH1 have been implicated separately as positive regulators of cancer cell migration and invasion in similar contexts (Arjonen et al., 2014; Carmona et al., 2016). In addition, MYO10 and RAPH1 knock-out mice share similar phenotypes, such as white belly patches due to defective melanoblast migration (Heimsath et al., 2017; Law et al., 2013). Therefore it is tempting to speculate that the MYO10-RAPH1 interaction occurring at filopodia tips has strong relevance in health and disease. Future work will investigate the contribution of the MYO10-RAPH1 interaction in regulating cell migration in vivo.

## Material and methods

### Cells

U2-OS osteosarcoma cells and MDA-MB-231 cells were grown in DMEM (Dulbecco’s Modified Eagle’s Medium; Sigma, D1152) supplemented with 10% fetal bovine serum (FCS) (Biowest, S1860). U2-OS cells were purchased from DSMZ (Leibniz Institute DSMZ-German Collection of Microorganisms and Cell Cultures, Braunschweig DE, ACC 785). MDA-MB-231 cells were provided by ATCC. The U2-OS MYO10-GFP lines were generated by transfecting U2-OS cells using lipofectamine 3000 (ThermoFisher Scientific), selected using Geneticin (ThermoFisher Scientific; 400 μg.ml-1 final concentration), and sorted for green fluorescence using a fluorescence-assisted cell sorter (FACS). All cell lines tested negative for mycoplasma.

### Antibodies and reagents

Mouse monoclonal antibodies used in this study were anti-β-actin (AC-15, Merck, A1978), anti-His tag (ThermoFisher Scientific, MA1-21315), and anti-tubulin (DHSB, clone 12G10). Rabbit polyclonal antibodies used in this study were anti-RAPH1 (ThermoFisher Scientific, PA5-110270), anti-MYO10 (Novus Biologicals, 22430002), and anti-GFP (Abcam, Ab290). Biotinylated proteins were detected using Streptavidin conjugated with Alexa Fluor™ 555 (for immunofluorescence) or Alexa Fluor™ 647 (for western blots), both provided by Thermo Fisher Scientific (S21381 and S21374). The bovine plasma fibronectin was provided by Merck (341631).

### Plasmids and transfection

U2-OS and MDA-MB-231 cells were transfected using Lipofectamine 3000 and the P3000TM Enhancer Reagent (Thermo Fisher Scientific) according to the manufacturer’s instructions.

The EGFPC1-hMyoX (MYO10-GFP) plasmid was a gift from Emanuel Strehler (Addgene plasmid 47608) (Bennett et al., 2007). The mScarlet-MYO10 (MYO10-RFP) construct was described previously (Jacquemet et al., 2019) and is available on Addgene (plasmid 145179). The mScarlet-I-MYO10^ΔF^ (MYO10^ΔF^-RFP) construct was previously described (Miihkinen et al., 2021) and is also available on Addgene (plasmid 145139). The GFP-VASP (mEmerald-VASP-N-10) plasmid was a gift from Michael Davidson (Addgene plasmid 54297). The GFP-RIAM(1-666) construct was a gift from Chinten James Lim (Addgene plasmid 80028) (Lee et al., 2013). The pcDNA3.1 MCS-BirA(R118G)-HA construct was a gift from Kyle Roux (Addgene plasmid 36047) (Roux et al., 2012). The RAPH1-GFP (EGFP-Lpd) plasmid was kindly provided by Matthias Krause (King’s College London).

The GFP-MYO10-BioID construct was generated as follows. Flanking XbaI sites were introduced into BioID by PCR (template plasmid: BioID pcDNA3.1 MCS-BirA(R118G)-HA) and the resulting amplicon was then inserted into a unique XbaI site in the EGFPC1-hMyoX plasmid resulting in an EGFP-MYO10-(stop codon)-BioID fusion gene. The stop codon between MYO10 and BioID was then replaced with a codon encoding valine (GTA) using a quick-change mutagenesis kit from Agilent and following the manufacturers’ instructions. The GFP-RAPH1^ΔTBS^ (RAPH1 aa 2-92 deleted) construct was created by inserting a custom gene block (IDT) in the EGFP-Lpd plasmid using the XhoI/HindIII sites. The RAPH1 fragments F1 (RAPH1 aa 1-535), F2 (RAPH1 aa 535-868), F3 (RAPH1 aa 535-868), and F4 (RAPH1 aa 868-1062) constructs were purchased from GenScript. Briefly, the gene fragments were synthesized using gene synthesis and cloned into pcDNA3.1(+)-N-eGFP using the BamHI/XhoI sites. The GST-RAPH1^F2^ construct (RAPH1 aa 535-868) was purchased from GenScript.

The gene fragment was synthesized using gene synthesis and cloned into pGEX-4T-1 using the BamHI/XhoI sites.

The GFP-MYO10-BioID, GFP-RAPH1^ΔTBS^, and RAPH1 fragments constructs will be made available on Addgene.

### siRNA-mediated gene silencing

The expression of RAPH1 was suppressed using 83 nM siRNA and lipofectamine 3000 (Thermo Fisher Scientific) according to the manufacturer’s instructions. siRNAs used were RAPH1 siRNA #2 (Hs_RAPH1_2, SI00698642) and RAPH1 siRNA #5 (Hs_RAPH1_5, SI04300982) provided by Qiagen.

### SDS–PAGE and quantitative western blotting

Protein extracts were separated under denaturing conditions by SDS–PAGE and transferred to nitrocellulose membranes using a Trans-Blot Turbo nitrocellulose transfer pack (Bio-Rad, 1704159). Membranes were blocked for 45 min at room temperature using 1x StartingBlock buffer (Thermo Fisher Scientific, 37578). After blocking, membranes were incubated overnight with the appropriate primary antibody (1:1000 in PBS), washed three times in TBST, and probed for 40 min using a fluorophore-conjugated secondary antibody diluted 1:5000 in the blocking buffer. Membranes were washed three times using TBST, over 15 min, and scanned using an Odyssey infrared imaging system (LI-COR Biosciences).

### GFP-trap pull-down

Cells transiently expressing bait GFP-tagged proteins were lysed in a buffer containing 20 mM HEPES, 75 mM NaCl, 2mM EDTA, 1% NP-40, as well as a cOmplete™ protease inhibitor tablet (Roche, cat. no. 5056489001), and a phosphatase inhibitor mix (Roche cat. no. 04906837001). Lysates were then centrifuged at 13,000 g for 10 min at 4C. Clarified lysates were incubated with GFP-Trap agarose beads for 2 h at 4C. Complexes bound to the beads were isolated by centrifugation, washed three times with ice-cold lysis buffer, and eluted in Laemmli reducing sample buffer.

### Protein expression and purification

The BL-21(DE3) E. coli strain was transformed with plasmids encoding the relevant His-tagged or GST-tagged proteins. Bacteria were grown at 37°C in LB media supplemented with ampicillin (100 μg/ml). Protein expression was induced with IPTG (1 mM) at 20°C. After 5 h, bacteria were harvested by centrifugation (20 min at 6000 g) and resuspended in resuspension buffer (1x TBS, Pierce Protease Inhibitor Tablet (Thermo Scientific, cat.no. A32963), 1X PMSF, 0.05mg/ml RNase, 0.05mg/ml DNase)). Bacteria were then lysed by adding BugBuster (Merck Millipore, cat. no. 70584-4) and a small spoonful of lysozyme (Thermo Scientific, cat. no. 89833). The suspension was mixed at 4°C for 30 min. Cell debris were then pelleted by ultracentrifugation (at 20000 rpm, JA25.50 rotor) at 4°C for 1 h. His-tagged MYO10 FERM was purified using a Protino Ni-TED 2000 packed column (Macherey Nagel, cat. no. 745120.25) according to the manufacturer’s instructions. The protein was eluted in multiple 1 ml fractions, supplemented with 1 mM AEBSF, and kept at 4°C until needed (up to one week). For GST-tagged proteins, 600 μl of equilibrated Glutathione Sepharose 4B beads (GE Healthcare, cat. no. 17-0756-01) were added to the supernatant and agitated for 1 h at 4°C. Beads were collected and washed four times with TBS supplemented with PMSF (1 mM). Protein-bound beads were stored at −80°C until needed.

### GST pull-down

GST and GST-RAPH1^F2^ sepharose beads were incubated with 10 mM His-tagged MYO10^FERM^, and the mixture was rotated overnight at 4°C. Beads were then washed four times with TBS supplemented with 1 mM PMSF. Proteins bound to beads were then eluted in 2x Laemmli sample buffer at 80°C. Results were then analyzed by western blot.

### Proximity biotinylation

U2-OS cells stably expressing GFP-MYO10 or GFP-MYO10-BioID were plated on fibronectin-coated plates in a medium containing 50 μM biotin for 24 h. After washing cells with cold PBS, cells were lysed, and debris were removed by centrifugation (13 000 x g, +4°C, 2 minutes). Biotinylated proteins were then incubated with streptavidin beads (MyOne Streptavidin C1, Invitrogen) for 1 h with rotation at +4°C. Beads were washed twice with 500 μl wash buffer 1 (10 % [w/v] SDS), once with 500 μl wash buffer 2 (0.1 % [w/v] deoxycholic acid, 1 % [w/v] Triton X-100, 1 mM EDTA, 500 mM NaCl, and 50 mM HEPES), and once with 500 μl wash buffer 3 (0.5 % [w/v] deoxycholic acid, 0.5 % [w/v] NP-40, 1 mM EDTA, and 10 mM Tris/HCl, pH 7.4). Proteins were eluted in 40 μl of 2 × reducing sample buffer for 10 min at 90°C.

### Mass spectrometry analysis

Affinity-captured proteins were separated by SDS-PAGE and allowed to migrate 10 mm into a 4–12% polyacrylamide gel. Following staining with InstantBlue (Expedeon), gel lanes were sliced into five 2-mm bands. The slices were washed using a solution of 50% 100 mM ammonium bicarbonate and 50% acetonitrile until all blue colors vanished. Gel slices were washed with 100% acetonitrile for 5–10 minutes and then rehydrated in a reducing buffer containing 20 mM dithiothreitol in 100 mM ammonium bicarbonate for 30 min at 56°C. Proteins in gel pieces were then alkylated by washing the slices with 100% acetonitrile for 5–10 minutes and rehydrated using an alkylating buffer of 55 mM iodoacetamide in 100 mM ammonium bicarbonate solution (covered from light, 20 min). Finally, gel pieces were washed with 100% acetonitrile, followed by washes with 100 μl 100 mM ammonium bicarbonate, after which slices were dehydrated using 100% acetonitrile and fully dried using a vacuum centrifuge. Trypsin (0.01 μg/μl) was used to digest the proteins (37°C overnight). After trypsinization, an equal amount of 100% acetonitrile was added, and gel pieces were further incubated at 37°C for 15 minutes, followed by peptide extraction using a buffer of 50% acetonitrile and 5% formic acid. The buffer with peptides was collected, and the sample was dried using a vacuum centrifuge. Dried peptides were stored at −20°C. Before LC-ESI-MS/MS analysis, dried peptides were dissolved in 0.1% formic acid. The LC-ESI-MS/MS analyses were performed on a nanoflow HPLC system (Easy-nLC1200, Thermo Fisher Scientific) coupled to the Orbitrap Fusion Lumos mass spectrometer (Thermo Fisher Scientific, Bremen, Germany) equipped with a nano-electrospray ionization source. Peptides were first loaded on a trapping column and subsequently separated inline on a 15 cm C18 column (75 μm x 15 cm, ReproSil-Pur 3 μm 120 Å C18-AQ, Dr Maisch HPLC GmbH, Ammerbuch-Entringen, Germany). The mobile phase consisted of water with 0.1% formic acid (solvent A) and acetonitrile/water (80:20 (v/v)) with 0.1% formic acid (solvent B). Peptides were eluted with 40 min method: from 8% to 43% of solvent B in 30 min, from 43% to 100% solvent B in 2 min, followed by a wash for 8 min at 100% of solvent B. MS data was acquired automatically by using Thermo Xcalibur 4.4 software (Thermo Fisher Scientific). A data-dependent acquisition method consisted of an Orbitrap MS survey scan of mass range 350-1750 m/z followed by HCD fragmentation for the most intense peptide ions in a full speed mode with a 2.5 sec cycle time.

Raw data from the mass spectrometer were submitted to the Mascot search engine using Proteome Discoverer 1.5 (Thermo Fisher Scientific). The search was performed against the human database SwissProt_2021_02, assuming the digestion enzyme trypsin, a maximum of two missed cleavages, an initial mass tolerance of 10 ppm (parts per million) for precursor ions, and a fragment ion mass tolerance of 0.020 Dalton. Cysteine carbamidomethylation was set as a fixed modification, and methionine oxidation was set as a variable modification.

To generate the MYO10-BioID dataset, five biological replicates were combined. Proteins enriched at least twofold in GFP-MYO10-BioID over GFP-MYO10 (based on spectral count) and detected with over five spectral counts (across all repeats) were considered putative MYO10 binder.

To generate the MYO10^FERM^ and Talin^FERM^ datasets, two biological replicates were combined. Proteins enriched at least twofold in MYO10^FERM^ over GFP and over Talin^FERM^ (based on spectral count) and detected with more than ten spectral counts (across both repeats) were considered putative MYO10 binders. The fold-change enrichment and the significance of the association used to generate the volcano Plots (Fig. 1A and 1B) were calculated directly in Proteome Discoverer.

### Light microscopy setup

The spinning-disk confocal microscope used was a Marianas spinning-disk imaging system with a Yokogawa CSU-W1 scanning unit on an inverted Zeiss Axio Observer Z1 microscope controlled by SlideBook 6 (Intelligent Imaging Innovations, Inc.). Images were acquired using either an Orca Flash 4 sCMOS camera (chip size 2,048 × 2,048; Hamamatsu Photonics) or an Evolve 512 EMCCD camera (chip size 512 × 512; Photometrics). The objective used was a 100x oil (NA 1.4 oil, Plan-Apochromat, M27) objective.

The structured illumination microscope (SIM) used was DeltaVision OMX v4 (GE Healthcare Life Sciences) fitted with a 60x Plan-Apochromat objective lens, 1.42 NA (immersion oil RI of 1.516) used in SIM illumination mode (five phases x three rotations). Emitted light was collected on a front-illuminated pco.edge sCMOS (pixel size 6.5 mm, read-out speed 95 MHz; PCO AG) controlled by SoftWorx.

The confocal microscope used was a laser scanning confocal microscope LSM880 (Zeiss) equipped with an Airyscan detector (Carl Zeiss) and a 40x water (NA 1.2) or 63x oil (NA 1.4) objective. The microscope was controlled using Zen Black (2.3), and the Airyscan was used in standard super-resolution mode.

### Quantification of filopodia numbers and dynamics

For the filopodia formation assays, cells were plated on fibronectin-coated glass-bottom dishes (MatTek Corporation) for 2 h.

Samples were fixed for 10 min using a solution of 4% PFA, then permeabilized using a solution of 0.25% (vol/vol) Triton X-100 for 3 min. Cells were then washed with PBS and quenched using a solution of 1 M glycine for 30 min. Samples were then washed three times in PBS and stored in PBS containing SiR-actin (100 nM; Cytoskeleton; catalog number: CY-SC001) at 4°C until imaging. Just before imaging, samples were washed three times in PBS. Images were acquired using a spinning-disk confocal microscope (100x objective). The number of filopodia per cell was manually scored using Fiji (Schindelin et al., 2012).

To study filopodia stability, U2-OS cells expressing MYO10-GFP were plated on fibronectin for at least 2 h before the start of live imaging (pictures taken every 5 s at 37°C, on an Airyscan microscope, using a 40x objective). All live-cell imaging experiments were performed in normal growth media, supplemented with 50 mM HEPES, at 37°C and in the presence of 5% CO_2_. Filopodia lifetimes were then measured by identifying and tracking all MYO10 spots using the Fiji plugin TrackMate (Tinevez et al., 2017; Ershov et al., 2022). In TrackMate, the custom Stardist detector and the simple LAP tracker (Linking max distance = 1 micron, Gapclosing max distance = 0 microns, Gap-closing max frame gap = 0 micron) were used. The StarDist 2D model used was trained for 200 epochs on 11 paired image patches (image dimensions: (512, 512), patch size: (512,512)) with a batch size of 2 and a mae loss function, using the StarDist 2D ZeroCostDL4Mic notebook (von Chamier et al., 2021; Schmidt et al., 2018). The training was accelerated using a Tesla K80 GPU.

### Generation of filopodia maps and analysis of filopodia length

U2-OS cells transiently expressing the constructs of interest were plated on high tolerance glass-bottom dishes (MatTek Corporation, coverslip 1.7) pre-coated first with Poly-L-lysine (10 μg/ml, 1 h at 37°C) and then with bovine plasma fibronectin (10 μg/ml, 2 h at 37°C). After 2 h, samples were fixed and permeabilized simultaneously using a solution of 4% (wt/vol) PFA and 0.25% (vol/vol) Triton X-100 for 10 min. Cells were then washed with PBS, quenched using a solution of 1 M glycine for 30 min, and, when appropriate, incubated with the primary antibody for 1 h (1:100). After three washes, cells were incubated with a secondary antibody for 1 h (1:100). Samples were then washed three times and incubated with SiR-actin (100 nM in PBS; Cytoskeleton; catalog number: CY-SC001) at 4°C until imaging (minimum length of staining, overnight at 4°C; maximum length, one week). Just before imaging, samples were washed three times in PBS and mounted in Vectashield (Vector Laboratories).

To map the localization of each protein within filopodia, images were first processed in Fiji (Schindelin et al., 2012), and data were analyzed using R as previously described (Jacquemet et al., 2019). Briefly, in Fiji, the brightness and contrast of each image were automatically adjusted using, as an upper maximum, the brightest cellular structure labeled in the field of view. In Fiji, line intensity profiles (1-pixel width) were manually drawn from filopodia tip to base (defined by the intersection of the filopodia and the lamel-lipodium). To avoid any bias in the analysis, the intensity profile lines were drawn from a merged image. All visible filopodia in each image were analyzed and exported for further analysis (export was performed using the “Multi Plot” function). For each staining, line intensity profiles were then compiled and analyzed in R. To homogenize filopodia length; each line intensity profile was binned into 40 bins (using the median value of pixels in each bin and the R function “tapply”). The map of each protein of interest was created by averaging hundreds of binned intensity profiles. The length of each filopodium analyzed was directly extracted from the line intensity profiles.

The preferential recruitment of protein to filopodia tips or shafts was assessed by calculating an enrichment ratio where the averaged intensity of the signal at the filopodia tip (bin 1-6) was divided by the averaged intensity at the filopodia shaft (bin 7-40).

### Quantification and statistical analysis

Randomization tests were performed using the online tool PlotsOfDifferences (https://huygens.science.uva.nl/PlotsOfDifferences/) (Goedhart, 2019). Dot plots were generated using PlotsOfData (Postma and Goedhart, 2019). Volcano Plots were generated using VolcaNoseR (Goedhart and Luijsterburg, 2020).

## Data availability

The authors declare that the data supporting the findings of this study are available within the article and from the authors on request.

## Acknowledgements

This study was supported by the Academy of Finland (G.J. 338537, and J.I, 325464; 346131), the Sigrid Juselius Foundation (J.I.), the Cancer Society of Finland (G.J. and J.I.), and Åbo Akademi University Research Foundation (G.J., CoE CellMech) and by Drug Discovery and Diagnostics strategic funding to Åbo Akademi University (G.J.). M.M. has been supported by the Drug Research Doctoral Programme, University of Turku foundation, Maud Kuistila foundation, Instrumentarium Foundation, Lounais-Suomen Syöpäyhdistys, K. Albin Johansson’s foundation, and Ida Montin foundation. B.T.G. was supported by the Biotechnology and Biological Sciences Research Council grant BB/S007245/1. A.P. was supported by the K. Albin Johanssons and the Swedish Cultural Foundations. We thank J. Siivonen and P. Laasola for technical assistance and H. Hamidi for editing the manuscript. The Cell Imaging and Cytometry Core facility (Turku Bioscience, University of Turku, Åbo Akademi University, and Biocenter Finland) and Turku Bioimaging are acknowledged for services, instrumentation, and expertise. Mass spectrometry analyses were performed at the Turku Proteomics Facility supported by Biocenter Finland.

## AUTHOR CONTRIBUTIONS

Conceptualization, G.J., and J.I.; Methodology, M.M., A.P., J.I. and G.J.; Formal Analysis, M.M., A.P., R.S., M.L.B.G. and G.J.; Investigation, M.M., A.P., S.G., R.S., M.L.B.G., N.J.B., B.T.G., J.I. and G.J.; Writing – Original Draft, G.J.; Writing – Review and Editing, M.M., A.P., S.G., R.S., M.L.B.G., B.T.G., J.I. and G.J.; Visualization, M.M., A.P., J.I. and G.J.; Supervision, G.J. and J.I.; Funding Acquisition, G.J. and J.I.

---

**Fig. S1.**
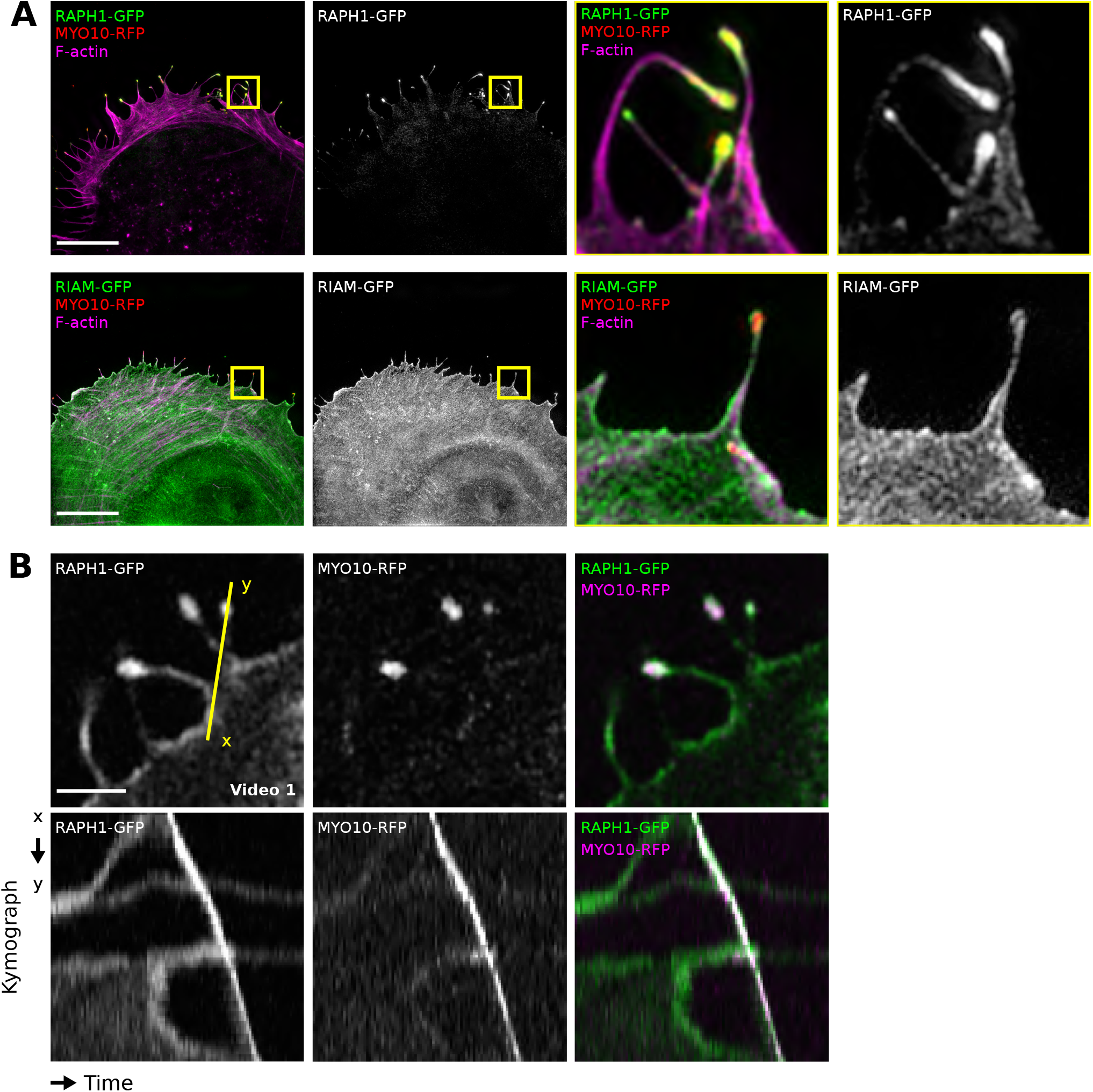
RAPH1, but not RIAM, accumulates at the tip of MYO10 filopodia. (**A**) U2-OS cells expressing MYO10-RFP and RAPH1-GFP or RIAM-GFP were plated on FN for 2 h, fixed, stained for F-actin, and imaged using structured illumination microscopy (SIM). Representative maximum intensity projections (MIPs) are displayed. The yellow squares highlight ROIs, which are magnified; scale bars: (main) 20 μm; (inset) 2 μm. (**B**) U2-OS cells expressing MYO10-RFP and RAPH1-GFP were imaged live at high spatiotemporal resolution using an Airyscan confocal microscope. A single time point (upper panel, scale bar: 2 μm) and a kymograph (lower panel) are displayed.

**Fig. S2.**
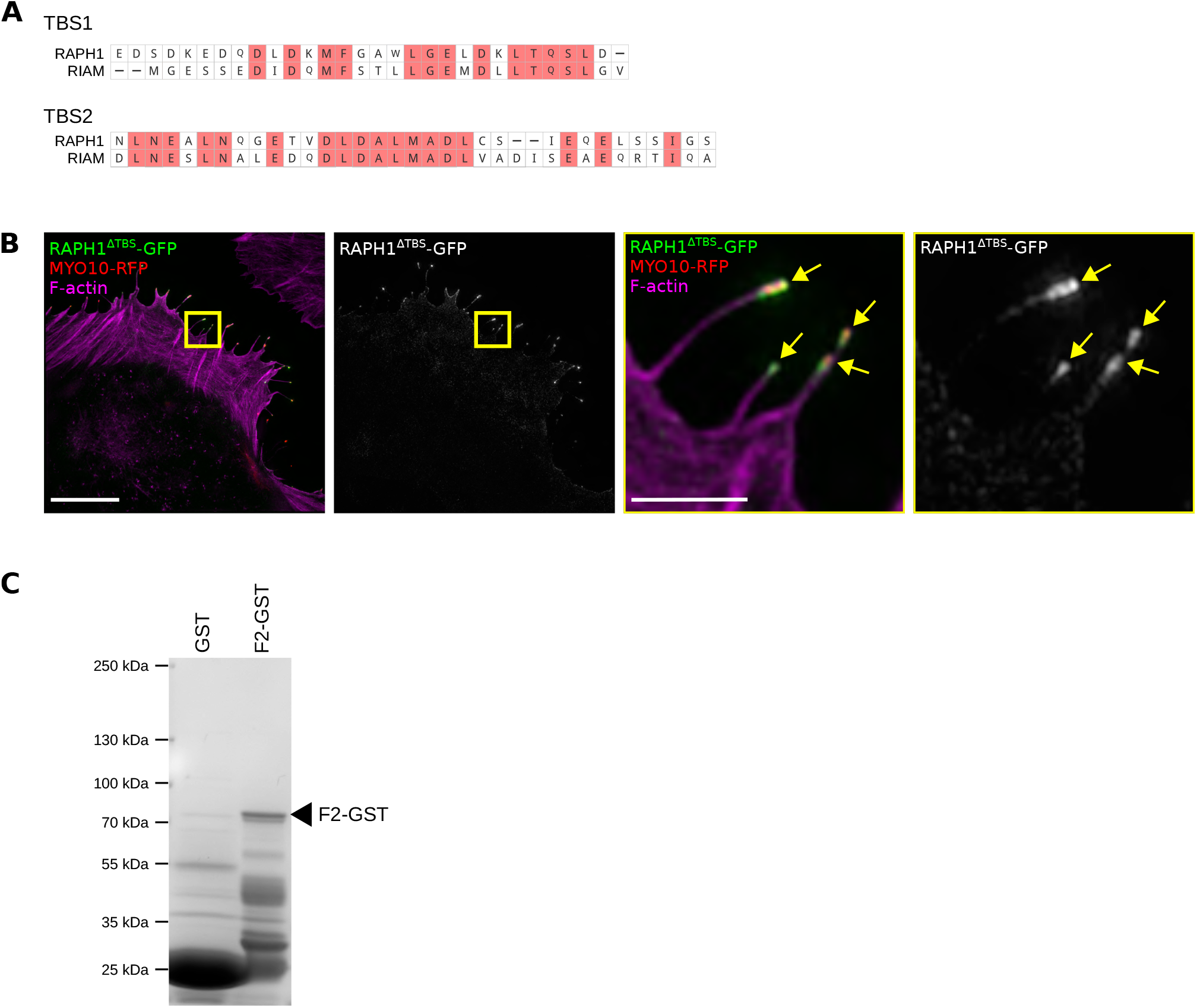
RAPH1 putative talin-binding sites do not contribute to its filopodia tip localization. (**A**) Alignment of the RAPH1 and RIAM talin-binding sites (TBS). (**B**) U2-OS cells expressing MYO10WT-RFP and GFP-RAPH1 lacking both TBS (GFP-RAPH1ΔTBS) were plated on FN for 2 h, fixed, stained for F-actin, and imaged using SIM. Representative MIPs are displayed. The yellow squares highlight ROIs, which are magnified; yellow arrows highlight filopodia tips; scale bars: (main) 10 μm; (inset) 2 μm. (**C**) Recombinant GST-RAPH1F2 and GST were produced in bacteria and subsequently purified using Glutathione agarose beads. Produced proteins were run on a polyacrylamide gel and stained using coomassie.

**Fig. S3.**
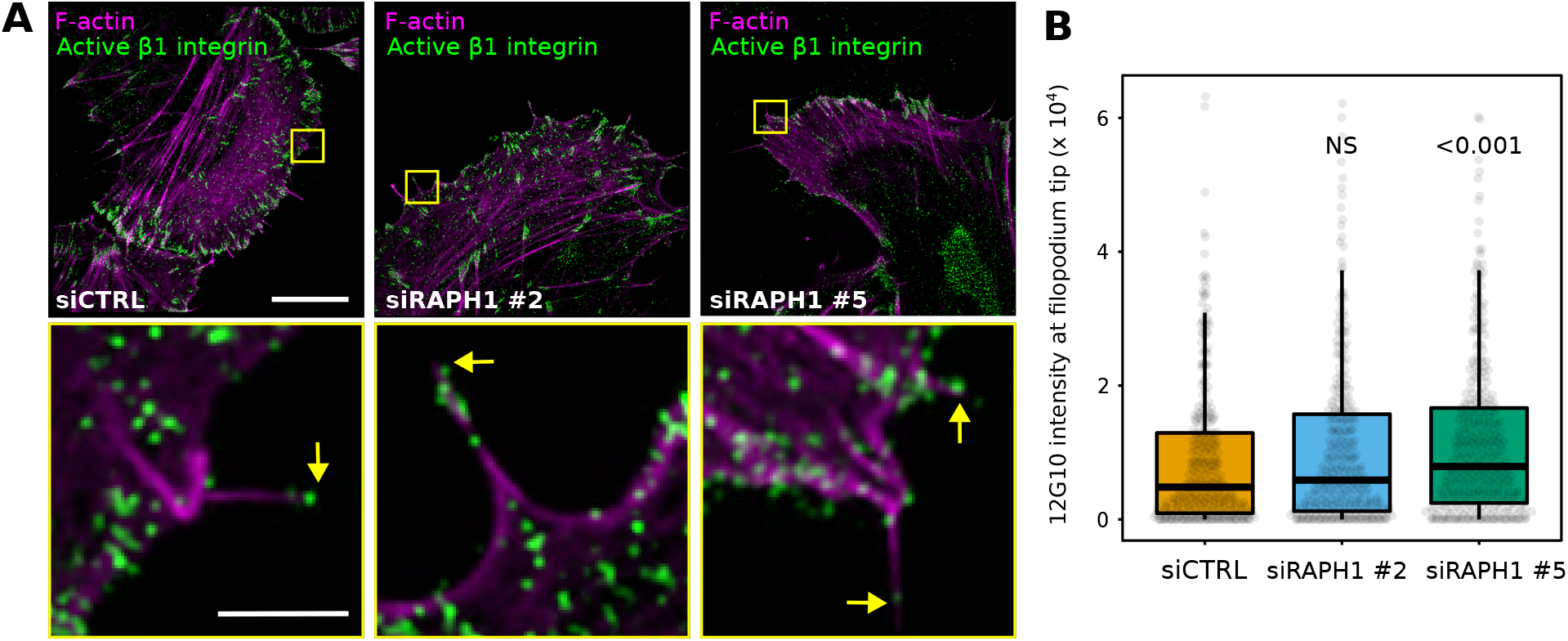
RAPH1 is not required for β1-integrin activation at filopodia tips. (**A**-**B**) RAPH1-silenced U2-OS cells expressing MYO10-GFP were plated on FN for 2 h, stained for active β1-integrin (12G10), and imaged using SIM. Representative MIPs are displayed; scale bars: (main) 20 μm; (inset) 2 μm. (**B**) The average intensity of 12G10 at filopodia tips was measured from line intensity profiles and displayed as boxplots (n > 400 filopodia, three biological repeats). P-values were determined using a randomization test. NS indicates no statistical difference between the mean values of the highlighted condition and the control.

